# Swainson’s thrushes vary song structure and singing behavior across ambient noise gradients and rapidly adjust songs in response to experimental traffic noise

**DOI:** 10.1101/2025.07.14.664765

**Authors:** Brigitta Mathews, Riley Crawford, Annika Dank, Jasper McCutcheon, Cvetana Ilic, Christopher N Templeton

**Affiliations:** Department of Biology, Western Washington University, Bellingham, WA 98225, USA; Cognitive Science, Pitzer College, Claremont, CA 91711, USA

## Abstract

Anthropogenic noise pollution has a variety of negative consequences for animals, with observed effects ranging from physiological processes to community structures. Sound waves from anthropogenic noise can mask animal signals, creating challenges for animal communication. Songbirds have been observed to combat noise pollution by changing their song rate, duration, frequency, or amplitude, with different species employing different strategies. We examined whether Swainson’s thrushes (*Catharus usulatus*) adjust their song structure under different traffic noise regimens using both population level surveys and individual level noise playback experiments. Swainson’s thrushes’ song frequency varied with background traffic noise levels across the surveyed population, but other song parameters did not. When confronted with realistic levels of experimental traffic noise playback, individual thrushes rapidly changed their song structure, increasing both the duration and the minimum frequency of their songs, before quickly returning towards baseline levels after the noise playback ended. Together, these results suggest that Swainson’s thrushes are adjusting their singing to reduce acoustic masking in noisy areas and further indicate that some songbirds might continually assess ambient noise levels and only adjust their signals when necessary.

## INTRODUCTION

Anthropogenic noise, noise produced by humans, has a wide variety of impacts on animals. While urban growth produces many types of pollution, anthropogenic noise is of particular importance due to its ability to spread beyond areas immediately impacted by urbanization to those without human development (Sordello et al., 2020). At the population and community levels, noise pollution can cause a reduction in biodiversity as usable habitats are altered and lost (Sordello et al., 2020). At the individual level, anthropogenic noise has negative implications for a variety of animal behaviors (Blackburn et al., 2024; Dowling et al., 2012; Gill et al., 2015; Hao et al., 2024; Laiolo, 2010), including reproductive success (Injaian et al., 2018, Mulholland et al., 2018), reduced predator defenses (Bruintjes & Radford, 2013), and changed foraging behaviors (Wang et al., 2022). Unsurprisingly, some of the largest behavioral impacts of noise pollution relate to auditory communication (Brumm, 2013).

Anthropogenic noise directly disrupts detection of auditory signals between both conspecifics and heterospecifics through auditory masking (Laiolo, 2010; Slabbekoorn & Ripmeester, 2008). For example, noise pollution disrupts the communication of anti-predator information in dwarf mongooses (*Helogale parvula*) through masking of surveillance calls (Kern & Radford 2016). Many species of marine mammals have also been observed to change their vocal signals in response to increasing levels of background noise. For example, Right whales (*Eubalaena glacialis*) increase their call amplitudes in noisy ocean environments to avoid masking (Parks et al., 2011) and killer whales (*Occinus orca*) increase the duration of their pod-specific social calls in response to increasing levels of boat traffic (Foote et al. 2004). Similar patterns are also seen in many other species, including monkeys, bats, and birds (Brumm & Zollinger, 2011).

In avian species, anthropogenic noise has been found to impact song structure in a variety of ways (Dowling et al., 2012; Wood & Yezerinac, 2006). Alterations in a bird’s song structure can have detrimental consequences for mating behaviors, territory defense, energy usage, and population density (Kuitunen et al., 2003; Slabbekoorn & Ripmeester, 2008; Collins, 2004; Luther et al., 2016). Consequently, understanding how anthropogenic noise impacts communication could help provide key insights on how noisy habitats impact avian fitness.

Traffic noise is particularly detrimental to avian communication because of its high-energy, low-frequency noise profile, which can mask the vocalizations of many songbirds (Derryberry et al., 2020). Species with lower natural vocalization frequencies are particularly vulnerable to the acoustic masking effect of traffic noise (Dowling et al., 2012; Francis et al., 2011; Nemeth et al., 2013). To compensate for acoustic masking from anthropogenic noise, birds have been shown to alter song structure by changing four different components of their vocalizations: amplitude, duration, song rate, and frequency (Brumm, 2013).

Changes in amplitude are primarily attributed to the Lombard Effect; first identified in 1911, this effect describes the phenomena of ‘speaking up’ in noisy environments (Brumm & Zollinger, 2011). An increase in vocal amplitude as a response to the presence of background noise has been identified in a variety of species including humans, monkeys, and whales (Brumm & Zollinger, 2011). This pattern is particularly prominent in birds. For example, a study of nightingales (*Luscinia megarhynchos*) across a natural ambient noise gradient revealed that birds in noisy habitats sang with a higher amplitude than those located in quieter habitats (Brumm, 2004). Similarly, great tits (*Parus major*) increased the amplitude of their alarm calls when faced with experimental noise in a controlled lab setting (Templeton et al., 2016).

Birds have also been shown to adjust song duration in the presence of noise. A study on great tits found that song lengths and inter-song intervals were shorter in city birds compared to their rural counterparts (Slabbekoorn & Den Boer-Visser, 2006). This could be a result of the non-forested urban habitat making it easier for songs to travel, causing birds to choose short songs to reduce the energetic cost of singing (Slabbekoorn & Den Boer-Visser, 2006). Alternatively, birds can avoid vocal masking by increasing signal redundancy, which can be achieved by increasing song duration (Brumm & Slater, 2006). A study on yellow warblers (*Setophaga petechia*) inhabiting different islands in the Galapagos found that on the island with lower human activity and less anthropogenic noise, the warblers decreased song duration in response to experimental noise playback. In contrast, on the noisier island, they increased song duration during noise exposure (Hohl et al., 2024). Similarly, a study on House finches (*Haemorhous mexicanus*) showed that duration increased with lower-frequency notes but decreased with higher-frequency notes in response to noise playback (Bermudez-Cuamatzin et al., 2011). Both studies support the idea that the relationship between duration and noise may depend on factors such as a bird’s prior exposure to certain noise levels and the frequency of their vocalizations.

Similar to song duration and amplitude, the rate at which birds vocalize is altered in noisy environments (LaZerte et al., 2017; Blackburn et al., 2024). Changes in song rate in response to increased anthropogenic noise have been observed in previous studies, but the changes are not consistent across species. A study comparing European robins (*Erithacus rubecula*) in noisy and quiet areas in Northern Ireland showed that birds inhabiting noisier areas sang fewer and less complex songs compared to those in quiet areas (Montague et al., 2012). In contrast, a study investigating zebra-finch response to growing noise levels revealed that they increase song rate up to an ambient noise threshold of 85 dB, at which point the rate begins to decrease or singing halts altogether (Brumm et al. 2013). These represent two different strategies that birds use to modulate song rate to promote communication during acoustic masking conditions. For some species, an increased song rate in noisier conditions may improve signaling by increasing time spent singing and therefore increasing signal redundancy (Brumm & Slater, 2006), while decreased song rate may be an attempt to avoid competition with low-frequency anthropogenic noise (Fuller, 2007).

Finally, anthropogenic noise has been shown to affect song frequency. Common changes in song frequencies include spectral shifts and omission of low-frequency notes (Slabbekoorn & Den Boer-Visser, 2006). Spectral shift refers to either a frequency change in all notes within a vocalization or of only the lowest frequency notes (Dowling et al., 2012; Hao et al., 2024; Slabbekoorn & Den Boer-Visser, 2006). Peak frequency, the frequency with the maximum amplitude in a signal, can vary if a bird sings a different song type or changes the spectral energy within the same song type without changing other features (Nemeth et al., 2013). Bandwidth, the frequency range between the minimum and maximum frequencies, is a useful marker for comparing spectral shifts and low note omission. The bandwidth of songs has been shown to decrease in noisy conditions, but this occurrence is not consistent across species and studies (Job et al., 2016; Rhodes et al., 2023). A common tactic for birds is to raise the minimum frequency of songs as ambient noise increases (Dowling et al., 2012; Derryberry et al., 2016; Slabbekoorn & Den Boer-Visser 2006; Wood & Yezerinac 2006). During the COVID-19 lockdown when anthropogenic traffic noise decreased, birds sang songs with lower minimum frequencies and increased bandwidth (Derryberry et al., 2020). This indicates that song frequency shifts can be plastic, potentially allowing birds to better communicate in adverse conditions.

In this study, we investigate the impact of anthropogenic noise on the song structure of Swainson’s thrush *(Catharus ustulatus*) songs. The Swainson’s thrush has a distinct, flute-like song that ascends in pitch as it progresses (Mack and Yong, 2000). The song begins with a low-frequency introductory note and is followed by 4-6 syllables that are longer, louder, and of higher frequency than the first note (Dobson and Lemon, 1977). Breeding in coniferous forests, Swainson’s thrushes are found in North America during spring and summer and migrate to Central and South America for the non-breeding season (Mack and Yong, 2000). During the breeding season, males sing regularly throughout the day to attract mates and to establish and defend territories. Swainson’s thrushes have a repertoire of 3-7 song types that are typically sung in a specific, fixed order (Dobson and Lemon, 1977). Because their breeding range overlaps with growing urban developments, Swainson’s Thrushes’ exposure to anthropogenic noise is likely increasing in parallel with increases in urbanization across western Washington. Here, we examine whether anthropogenic noise impacts Swainson’s thrush song structure and singing behavior to determine the potential short and long-term impacts of noise pollution and strategies that this species uses to combat acoustic masking from noise pollution.

Following a protocol used by LaZerte et al. (2017), we evaluated Swainson’s thrush song structure and behavior by combining both population level recordings of birds singing across a gradient of background noise levels and conducting experimental anthropogenic noise playbacks to identify rapid song structure changes in individuals. To determine which strategies Swainson’s thrushes are employing, if any, we analyzed different song features, including song duration, song bandwidth, song rate, and three measures of song frequency. We predicted that Swainson’s thrushes would attempt to avoid masking effects of low-frequency traffic noise by increasing minimum and peak frequencies and reducing the duration of songs as ambient noise levels increase, in both observational and experimental situations. We also predicted that due to its low frequency, the introductory note of the Swainson’s thrush song would show more pronounced effects as it is likely most susceptible to masking.

## METHODS

### Study Sites

We studied Swainson’s thrushes in locations within and nearby Bellingham, Washington between June 26 and August 8, 2024. 43 individual vocalizing males were recorded in urban parks, green spaces, and forests located in areas with varying levels of traffic noise (Figure 1). Sites ranged from immediately adjacent to a busy interstate highway (I-5) to pristine forest locations without significant noise pollution, with specific sites chosen to capture a gradient of ambient anthropogenic noise across thrush breeding territories. To ensure the same individuals were not recorded more than once, we selected sites that were at least 400 meters apart. Ambient noise levels of our study sites ranged from 30.5 dB to 73.3 dB.

**Figure 1.**
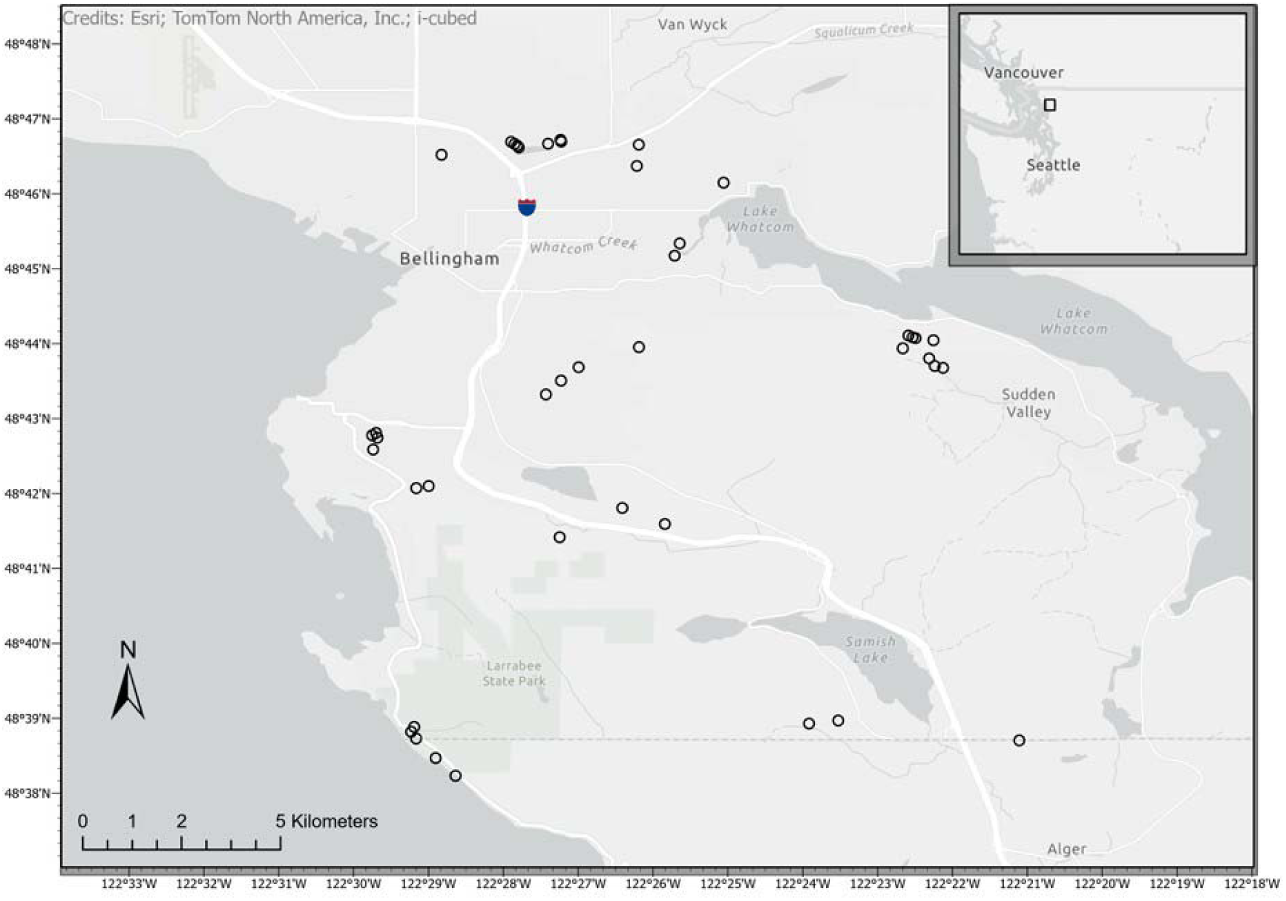
Study sites across Western Washington, showing locations of study sites (points), and major sources of traffic noise (white lines), including I5 (freeway marker) and other highways.

**Figure 2.**
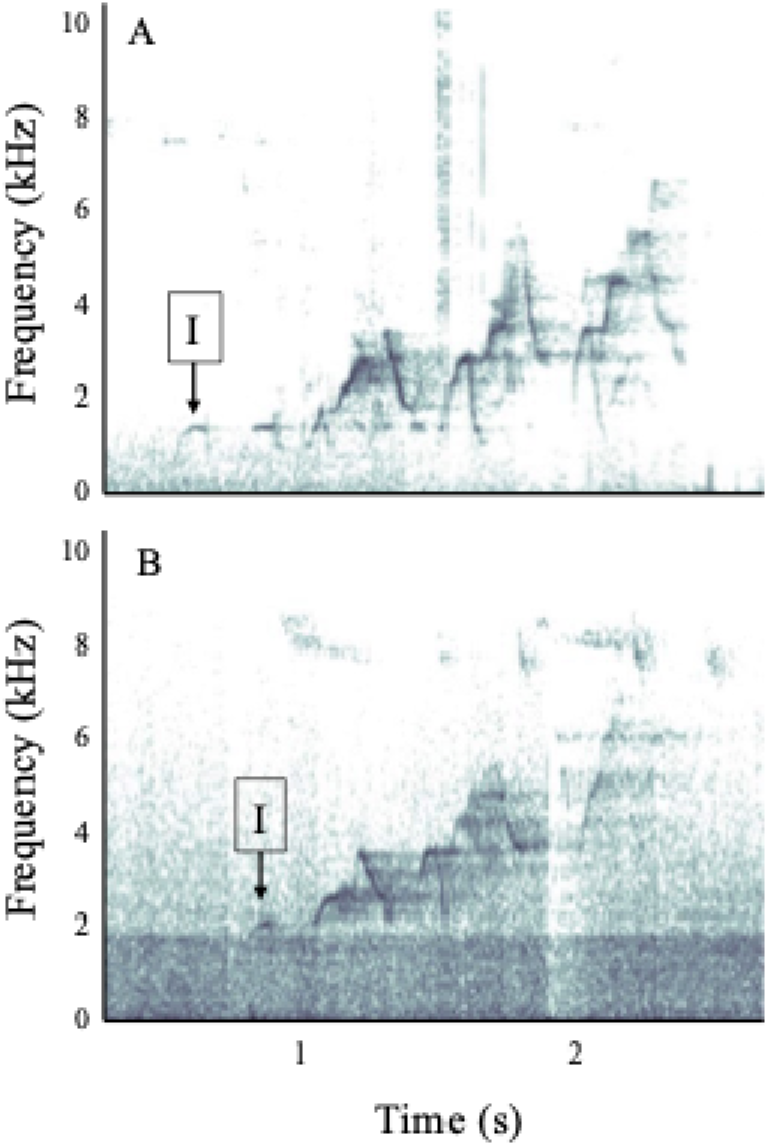
Swainson’s thrushes sing an ascending and melodious song. Songs were recorded in both (A) relatively quiet locations and those with (B) higher levels of background traffic noise. Song features were measured both for the entire song and for the introductory syllable, designated with an I.

### Ambient Noise Measurements

We measured ambient noise levels at each site by using an iPhone with the NIOSH Sound Level Meter app (settings were calibrated to Threshold Level = 80, Exchange Rate = 3dB, Time Weighting = Slow, Frequency weighting = A). We made two 60-second recordings 1.5 meters above the ground and took the average LAeq (dB) reading to account for natural variation in ambient noise. The first measurement was taken during the beginning of the survey period, and the second measurement was taken towards the end of the survey. For birds that were exposed to experimental playback, the first measurement was taken in the pre-playback period and the second was taken in the post-playback period. For 8 of the 43 birds, only one noise recording was taken, which was used in place of an average.

### Population Level Song Survey

To assess how song parameters vary with background noise, we recorded birds across a traffic noise gradient. Recordings were taken of singing male Swainson’s thrushes using Marantz PMD661 recording devices and Sennheiser ME66/K6 shotgun mics approximately10 meters from the focal individual. Recorders were set to record uncompressed .wav files at a 44kHz sampling rate. We recorded the focal individual for five minutes. We completed surveys with 42 individual birds.

### Individual Responses to Noise Playback

To experimentally assess how individuals respond to varying traffic noise levels, we conducted a repeated measures playback experiment with individual Swainson’s thrushes. If a bird was still vocalizing at the end of the five-minute survey period, we then exposed it to a five-minute recording of traffic noise. Experimental traffic noise was recorded from a moderately busy highway and provided realistic levels of highway noise (Templeton et al., 2016; Osbrink et al., 2021). We played this traffic noise from a calibrated FoxPro Crossfire speaker to add approximately 65 dB of anthropogenic noise during playback (LaZerte et al., 2017) when the speaker was placed ca. 10m from the focal bird at 1-2m height. We randomly selected one of five different exemplar noise recordings to play at each site using a random number generator. After the noise playback finished, we continued recording for five minutes post-playback. We conducted experimental trials with 15 individuals.

### Acoustic Analysis

Analysis of the high-quality audio recordings were performed using Raven Pro v.1.5 software (Cornell University, Ithaca NY USA) to measure our chosen variables for each individual Swainson’s thrush. The Raven selection tool was used to visually select and box up to 10 (minimum = 3, average = 8) full songs from each survey recording and up to 10 from each of the playback periods for experimental trials (pre playback, during noise playback, and post). Some recordings did not have 10 high quality songs available for selection due to overlap of other vocalizations, loud background noises, or poor audio quality. Because of their relatively low frequency, we also separately identified and boxed the introductory note of each song. Boxing bias was reduced by following a uniform protocol (Charif et al., 2010), with measurements taken from spectrograms with the following parameters: window type: Hann, window size: 1024 Hz, 3 dB filter bandwidth: 67.4 Hz, brightness: 75, contrast: 65. We used Raven Pro to automatically calculate acoustic parameters that are robust to variation in boxing ( Charif et al., 2010 , and focused the analyses on Duration 90% (s), Peak Frequency Contour (PFC) Max Frequency (Hz), PFC Min Frequency (Hz), Peak Frequency (Hz), and calculated song Bandwidth as the difference between PFC Max and Min frequencies. Vocalizations boxed from the pre-playback segment of the experimental trials were also used as survey data points for applicable individuals, as the birds vocalizing in this phase of the experiment had not yet been exposed to experimental manipulations. To calculate song rate, every individual song was counted from the pre-, during-, and post-playback periods of each experimental noise trial and we calculated the average songs per minute for each trial period.

### Statistical Analyses

To assess the impact of anthropogenic noise on Swainson’s thrush song frequency, duration, and rate, we created and ran linear mixed effect models (LMMs) using the lme4 R package in R v.2024.12.0+467 (R Core Team 2025; Bates et al. 2015). We fit LMMs using restricted maximum likelihood (REML) as this approach performs better than maximum likelihood with small sample sizes (Luke 2016). We used a combination of Shapiro-Wilk W tests, Levene’s tests, and visual inspections of diagnostic plots to assess whether the model assumptions were met (Jacqmin-Gadda et al. 2007), including normality of the random effect, normality of residuals, residual homoscedasticity, and the independency of the residuals. The diagnostic plots we created to decide whether our models met the assumptions were a residuals versus fitted values plot, a QQ plot for normality of residuals, a QQ plot for normality of random effects, and an autocorrelation function (acf) plot.

We extracted p-values using the lmerTest package (Kuznetsova et al. 2017) to assist in determining the statistical significance of the models. The critical value for all comparisons was 0.05 using two-tailed tests. The lmerTest package utilizes t-tests with Satterthwaite’s approximation to provide p-values for LMMs (Kuznetsova et al. 2017). We chose this method because of its applicability to REML models and reliability at assessing significance in LMMs (Luke 2016). In addition, we calculated the R^2^ values using the Performance package (Ludecke et al. 2021) for all statistically significant models. This allowed us to draw conclusions about the degree of biological significance of our results. This package calculates the marginal R^2^ values with the approach from Nakagawa et al. (2017), which we used to assess the contributions of each fixed effect.

### Population level song survey

We ran a LMM for each combination of vocalization type (full song or intro note) and response variable (minimum frequency, maximum frequency, bandwidth, peak frequency, and duration), which resulted in 10 unique models. The predictor variable was always ambient noise level (dB). We recorded songs from 42 unique individuals; for the models with the full song vocalization type we included 328 songs, while the models with the intro note type included 357 notes. To remove intra-individual variation from our data, we set individual ID (a unique identifier for each bird) as a random effect. Of our 10 total models, all but two met the assumptions of LMMs. The two problematic models failed to meet the residual homoscedasticity assumption. Because the heteroscedasticity in those models was not dramatic, and linear mixed models are generally robust to violations of this assumption (Jacqmin-Gadda et al. 2007; Schielzeth et al. 2020), we felt it acceptable to move forward with those models.

### Individual responses to noise playback

We also used LMMs to analyze the experimental traffic noise playback data. There were 11 models in total, each representing a unique combination of a vocalization type (full song or intro note) and a response variable (minimum frequency, maximum frequency, bandwidth, peak frequency, duration, and song rate). The predictor variable for all these models was playback phase (pre, noise, or post playback). We recorded songs from 15 unique individuals in each phase of the study; models with the full song vocalization type had a sample size of 119 pre, 62 during noise, and 90 post playback songs, while the intro note type had 126 pre, 64 during noise, and 90 post playback songs. For this suite of models, we also removed intra-individual variation from our data by using individual ID as a random effect. We concluded that all 11 models met the assumptions of LMMs following a visual inspection of the diagnostic plots of each.

To quantify how Swainson’s thrushes varied their song characteristics during the different portions of traffic noise playback, we calculated estimated marginal means (EMMs). These values represented how much the vocalizations of Swainson’s thrushes changed in response to playback. Here, EMMs indicate the predicted average values of the response variables after accounting for individual variation and were calculated using the emmeans R package (Lenth 2025).

## RESULTS

### Population level song survey

Swainson’s thrushes’ song features varied along a traffic noise gradient (Table 1). Survey data showed that Swainson’s thrushes significantly increased the minimum frequency of their songs as noise levels increased (p<0.001; Figure 3A). In contrast, the bandwidth (p = 0.004) and maximum frequency (p = 0.026) of their songs was lower as ambient noise increased (Figure 3B and C). The peak frequency and duration of Swainson’s thrushes’ songs were unaffected by ambient noise levels (p = 0.766, Figure 3D; p = 0.502, Figure 3E; respectively). Background noise moderately explained the remaining variation for minimum frequency (marginal R^2^ = 0.187) while poorly explaining that for maximum frequency, bandwidth, peak frequency, and duration (R^2^ = 0.053, 0.092, 0.001, 0.004, respectively).

**Figure 3.**
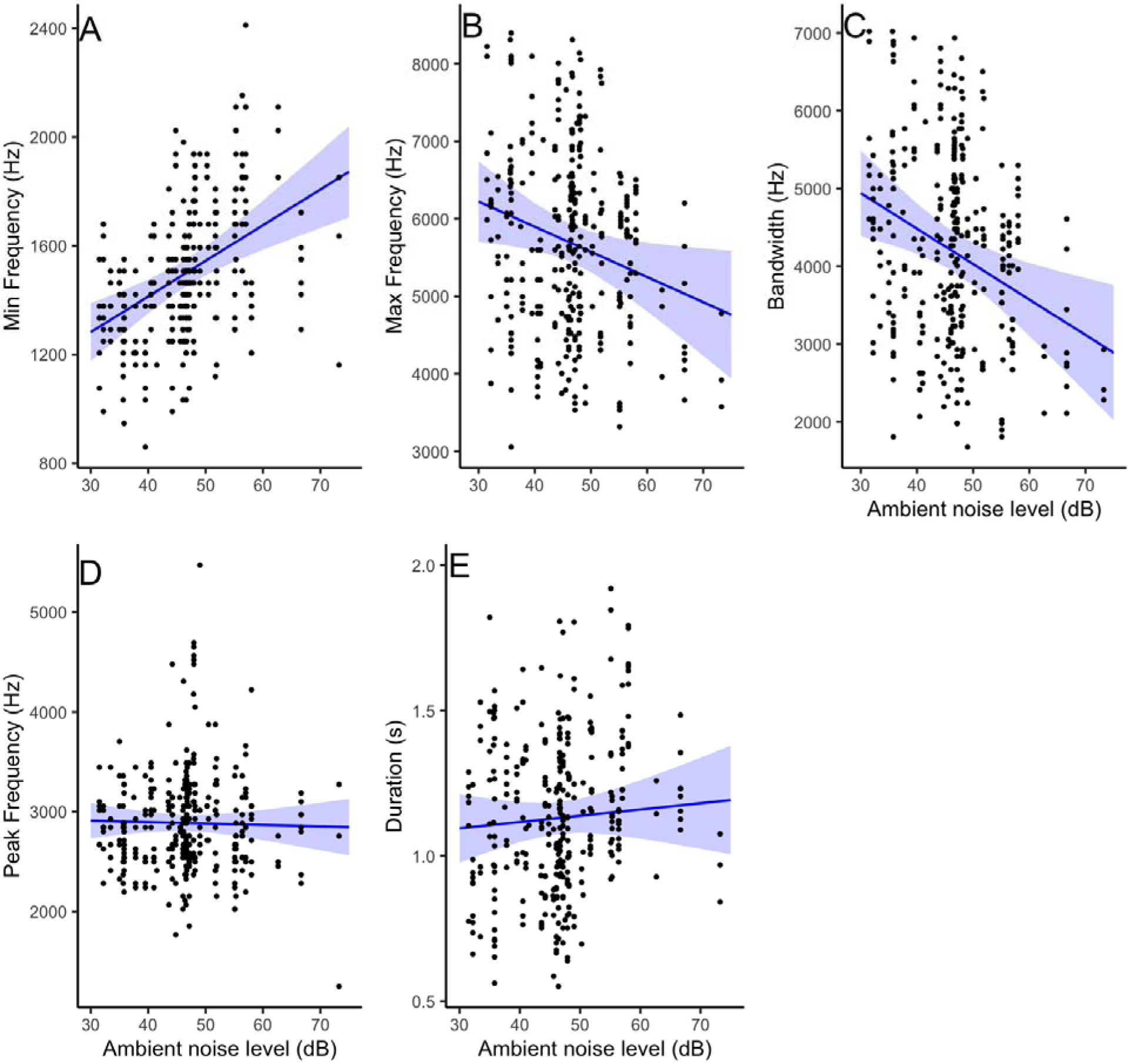
In areas with greater ambient noise levels, Swainson’s thrushes sang songs that had higher (A) minimum frequencies, lower (B) maximum frequencies, and shorter (C) bandwidths while their (D) peak frequency and (E) duration showed no patterns. Points represent raw data while the blue line and shaded areas indicate LMM 95% confidence intervals.

**Table 1.** Linear mixed effect model results for the impact of ambient noise on Swainson’s thrushes’ full song and intro note features. Columns for degrees of freedom (df), the t value, and P value were derived from t-tests with Satterthwaite’s approximation. Bold text indicates significant effects.

| Response Variable | Parameter | Estimate | Std. Error | df | t value | P value |
| --- | --- | --- | --- | --- | --- | --- |
| Full song minimum frequency (Hz) | Intercept | 889.62 | 136.50 | 42 | 6.52 | <0.001 |
|  | **Ambient noise (dB)** | **13.11** | **2.88** | **43** | **4.56** | **<0.001** |
| Full song maximum frequency (Hz) | Intercept | 7195.81 | 668.16 | 46 | 10.77 | <0.001 |
|  | **Ambient noise (dB)** | **-32.45** | **14.09** | **47** | **-2.30** | **0.026** |
| Full song bandwidth (Hz) | Intercept | 6306.10 | 708.31 | 46 | 8.90 | <0.001 |
|  | **Ambient noise (dB)** | **-45.56** | **14.93** | **46** | **-3.05** | **0.004** |
| Full song peak frequency (Hz) | Intercept | 2954.17 | 228.96 | 48 | 12.90 | <0.001 |
|  | Ambient noise (dB) | -1.45 | 4.84 | 49 | -0.30 | 0.766 |
| Full song duration (s) | Intercept | 1.03 | 0.15 | 44 | 6.80 | <0.001 |
|  | Ambient noise (dB) | 0.00 | 0.00 | 45 | 0.68 | 0.502 |
| Intro note minimum frequency (Hz) | Intercept | 1035.69 | 137.02 | 50 | 7.56 | <0.001 |
|  | **Ambient noise (dB)** | **11.77** | **2.88** | **51** | **4.08** | **<0.001** |
| Intro note maximum frequency (Hz) | Intercept | 1474.92 | 173.75 | 46 | 8.49 | <0.001 |
|  | **Ambient noise (dB)** | **12.40** | **3.66** | **47** | **3.39** | **0.001** |
| Intro note bandwidth (Hz) | Intercept | 419.42 | 126.91 | 46 | 3.30 | 0.002 |
|  | Ambient noise (dB) | 1.02 | 2.68 | 47 | 0.38 | 0.705 |
| Intro note peak frequency (Hz) | Intercept | 1366.64 | 157.02 | 46 | 8.70 | <0.001 |
|  | **Ambient noise (dB)** | **12.31** | **3.31** | **47** | **3.72** | **<0.001** |
| Intro note duration (s) | Intercept | 0.09 | 0.02 | 47 | 5.19 | <0.001 |
|  | Ambient noise (dB) | 0.00 | 0.00 | 48 | 0.68 | 0.503 |

Background noise levels had a strong effect on the structure of the low frequency introductory note of Swainson’s thrushes’ songs (Table 1). Analysis of intro notes showed that the minimum frequency (11.77 ± 2.88 Hz per dB; p<0.001; Figure 4A) increased comparably with the maximum frequency (12.40 ± 3.66 Hz per one dB increase; p = 0.001; Figure 4B).

**Figure 4.**
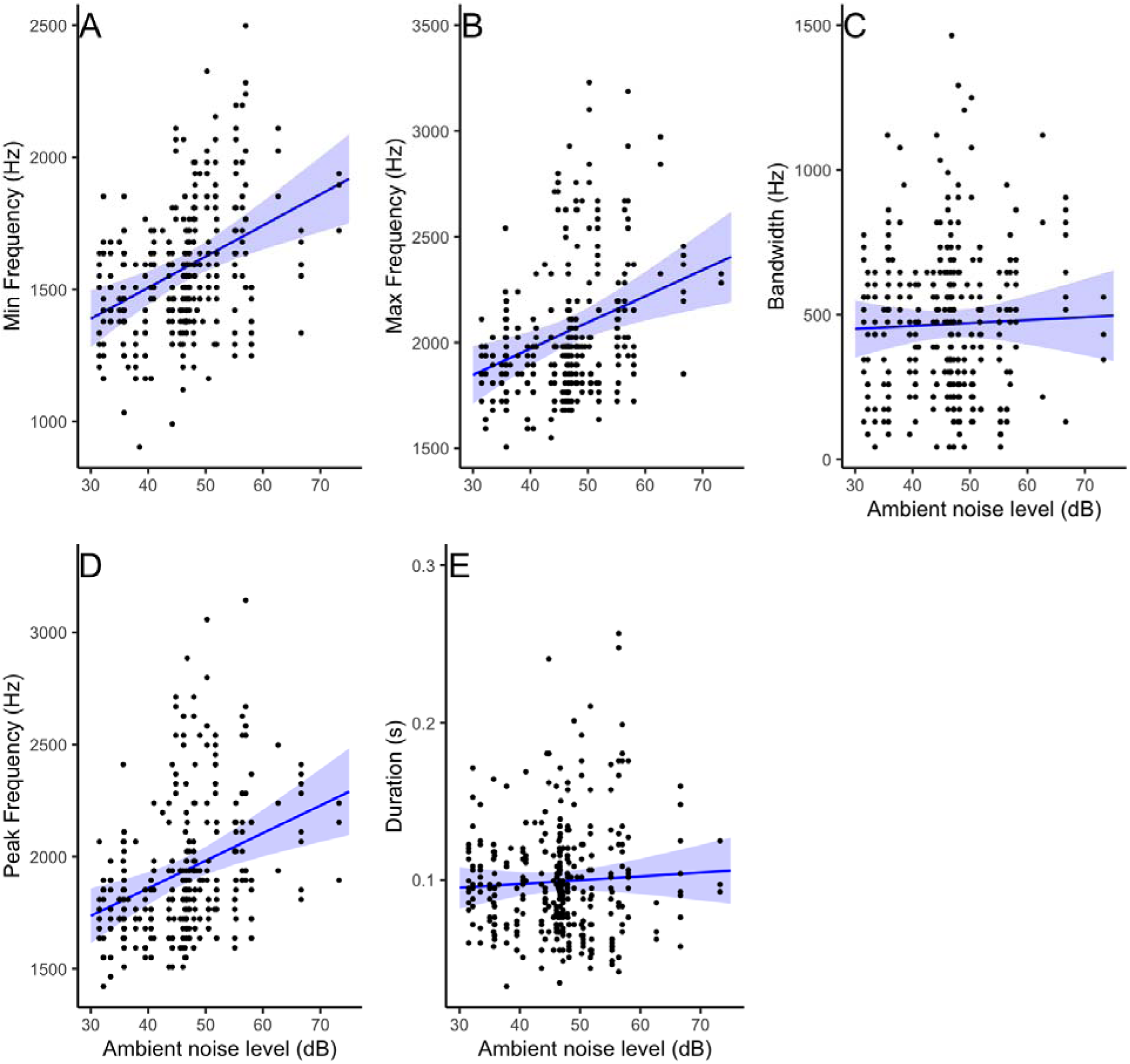
In areas with greater ambient noise levels, Swainson’s thrushes’ songs had intro notes with higher (A) minimum, (B) maximum, and (D) peak frequencies while their (C) bandwidth and (E) duration showed no patterns. Points represent raw data with the blue line and shaded areas indicating the estimate and 95% confidence intervals from the respective LMMs.

Similarly, the peak frequency of the intro notes was higher as background noise increased (p<0.001; Figure 4D). The bandwidth (p = 0.705; Figure 4C) and duration (p = 0.503; Figure 4E) of intro notes did not demonstrate any significant relationship with background noise levels. The variation seen in the minimum, maximum, and peak frequency was moderately explained by ambient noise (marginal R^2^ = 0.155, 0.106, and 0.115, respectively), which only poorly explained variation in bandwidth (R^2^ = 0.001) and duration (R^2^ = 0.003).

### Individual responses to noise playback

Individual Swainson’s thrushes rapidly changed the acoustic parameters of their songs when confronted with the experimental traffic noise playback (Table 2). The minimum frequency of Swainson’s thrushes’ songs significantly increased during noise playback (p<0.001) and remained statistically higher after the noise was played (p = 0.001), although a trend of reducing back towards baseline levels exists (Figure 5A). While song bandwidth was significantly reduced when exposed to noise playback, it returned to pre-playback levels once playback stopped (noise, p<0.001; post, p = 0.399; Figure 5C). Swainson’s thrushes increased the length of their songs in response to noise playback, with song length also returning to pre-playback levels after the noise playback ended (noise, p<0.001; post, p = 0.931; Figure 5E).

**Figure 5.**
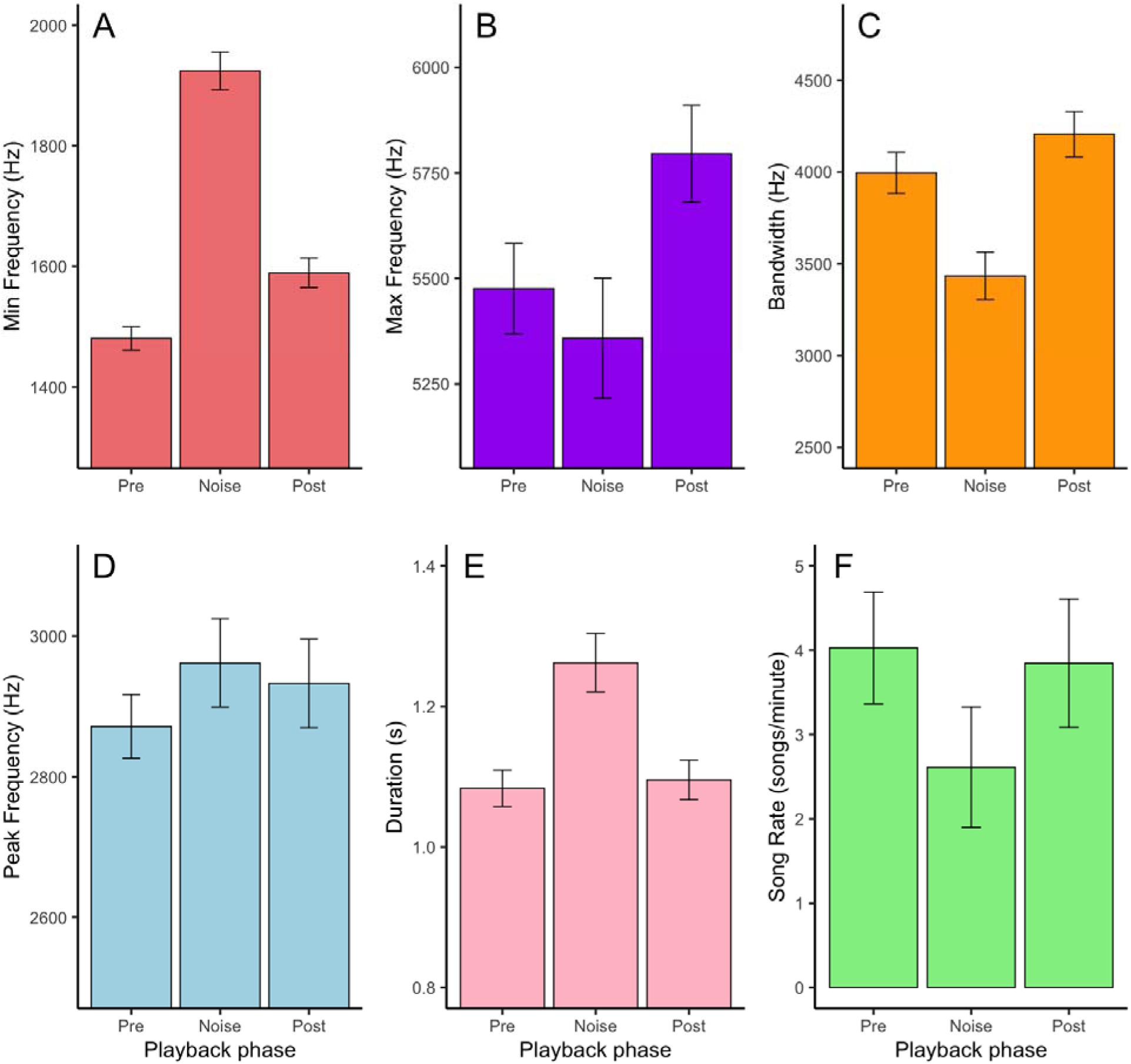
Swainson’s thrushes rapidly changed the minimum frequency, bandwidth, and duration of their songs upon exposure to the experimental traffic noise playback. Mean ± standard error are shown for the (A) minimum frequency, (B) maximum frequency (C) bandwidth, (D) peak frequency, (E) duration, and (F) song rate of Swainson’s thrushes’ full songs during the three playback phases (pre, noise, and post).

**Table 2.** Estimated marginal means results from a linear mixed effect model for the impact of experimental noise playback on Swainson’s thrushes’ full song and intro note features. Columns for degrees of freedom (df), the t value, and P value were all derived from t-tests with Satterthwaite’s approximation. The t-tests compared the during and post parameters/phases to the reference level (pre). Bold text indicates significant effects.

| Response Variable | Parameter | Estimate | Std. Error | df | t value | P value |
| --- | --- | --- | --- | --- | --- | --- |
| Full song minimum frequency (Hz) | Pre (intercept) | 1,465.64 | 40.67 | 16 | 36.04 | <0.001 |
|  | **Noise** | **1,892.30** | **44.92** | **263** | **13.93** | **<0.001** |
|  | **Post** | **1,555.95** | **42.47** | **265** | **3.30** | **0.001** |
| Full song maximum frequency (Hz) | Pre (intercept) | 5,557.95 | 196.88 | 16 | 28.23 | <0.001 |
|  | Noise | 5,271.17 | 222.31 | 265 | -1.73 | 0.085 |
|  | Post | 5,777.28 | 207.65 | 267 | 1.48 | 0.14 |
| Full song bandwidth (Hz) | Pre (intercept) | 4,090.86 | 200.31 | 16 | 20.42 | <0.001 |
|  | **Noise** | **3,378.44** | **227.13** | **265** | **-4.14** | **<0.001** |
|  | Post | 4,220.66 | 211.67 | 267 | 0.84 | 0.399 |
| Full song peak frequency (Hz) | Pre (intercept) | 2,877.12 | 60.86 | 27 | 47.28 | <0.001 |
|  | Noise | 2,973.27 | 79.32 | 261 | 1.14 | 0.257 |
|  | Post | 2,925.59 | 68.51 | 260 | 0.65 | 0.519 |
| Full song duration (s) | Pre (intercept) | 1.10 | 0.05 | 16 | 22.07 | <0.001 |
|  | **Noise** | **1.28** | **0.06** | **265** | **4.25** | **<0.001** |
|  | Post | 1.10 | 0.05 | 267 | -0.09 | 0.931 |
| Full song rate (songs/minute) | Pre (intercept) | 4.03 | 0.71 | 40 | 5.64 | <0.001 |
|  | Noise | 2.61 | 0.71 | 28 | -1.52 | 0.14 |
|  | Post | 3.85 | 0.71 | 28 | -0.19 | 0.848 |
| Intro note minimum frequency (Hz) | Pre (intercept) | 1,568.60 | 38.78 | 16 | 40.45 | <0.001 |
|  | **Noise** | **1,959.06** | **42.94** | **270** | **13.55** | **<0.001** |
|  | **Post** | **1,680.49** | **40.71** | **275** | **4.19** | **<0.001** |
| Intro note maximum frequency (Hz) | Pre (intercept) | 1,948.56 | 47.08 | 17 | 41.39 | <0.001 |
|  | **Noise** | **2,370.87** | **53.64** | **272** | **10.47** | **<0.001** |
|  | **Post** | **2,055.67** | **50.13** | **276** | **2.88** | **0.004** |
| Intro note bandwidth (Hz) | Pre (intercept) | 380.43 | 28.54 | 21 | 13.33 | <0.001 |
|  | Noise | 409.20 | 34.69 | 276 | 0.92 | 0.359 |
|  | Post | 372.75 | 31.41 | 276 | -0.27 | 0.789 |
| Intro note peak frequency (Hz) | Pre (intercept) | 1,854.43 | 46.45 | 17 | 39.92 | <0.001 |
|  | **Noise** | **2,273.16** | **52.83** | **272** | **10.62** | **<0.001** |
|  | **Post** | **1,944.04** | **49.42** | **276** | **2.46** | **0.014** |
| Intro note duration (s) | Pre (intercept) | 0.09 | 0.00 | 23 | 20.73 | <0.001 |
|  | Noise | 0.10 | 0.01 | 276 | 1.96 | 0.051 |
|  | Post | 0.09 | 0.01 | 275 | -0.07 | 0.944 |

Maximum frequency, peak frequency, and song rate did not significantly change between the different playback phases (Figures 5B, D, and F). The playback of traffic noise explained a meaningful proportion of the variation in the minimum frequency (marginal R^2^ = 0.355) of full songs while poorly explaining that of maximum frequency, bandwidth, peak frequency, duration, and song rate (R^2^ = 0.025, 0.069, 0.005, 0.061, 0.050, respectively). While there was some variation in responses across different individuals, most individuals responded to noise playback in a similar fashion (Figure 6).

**Figure 6.**
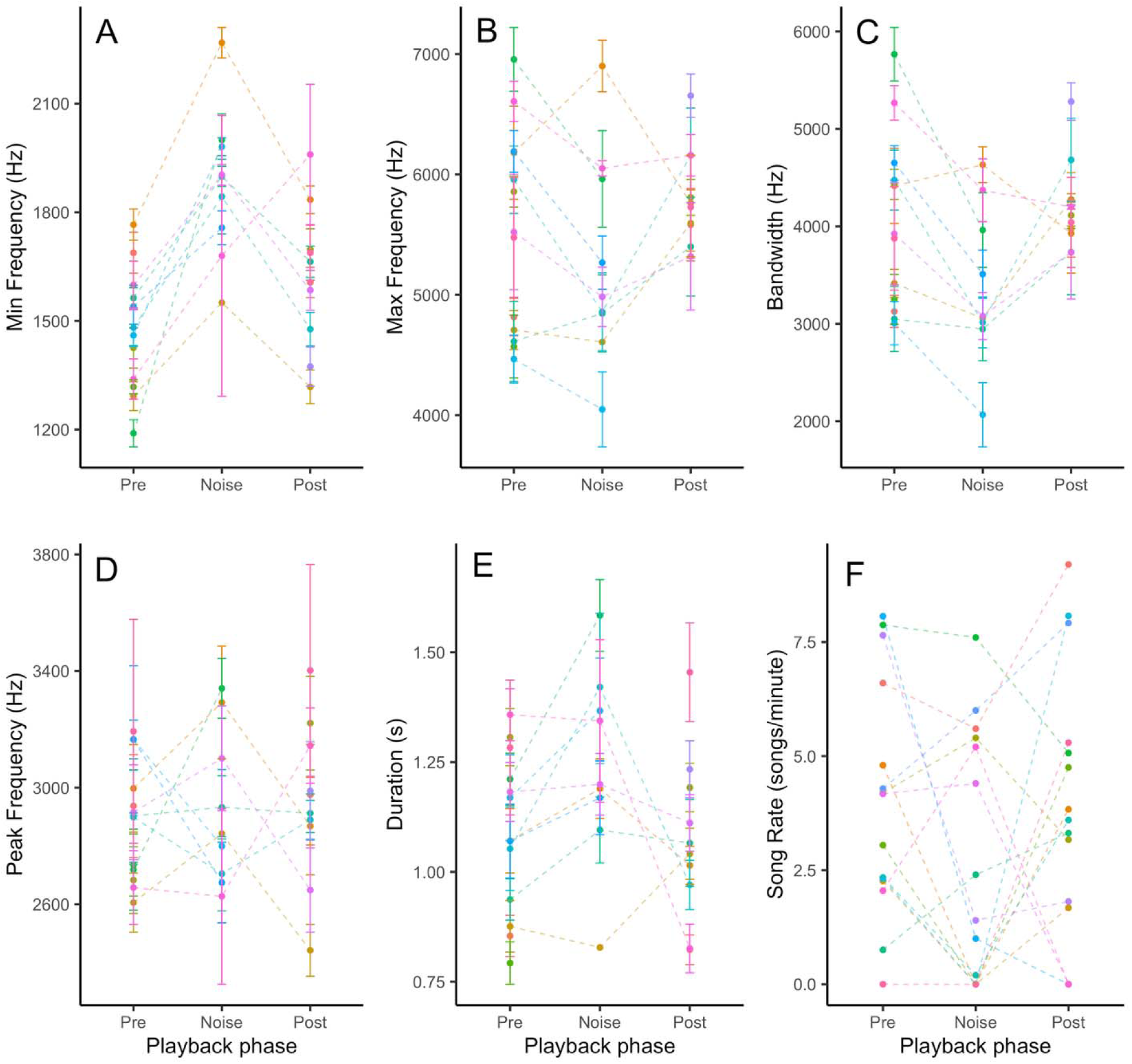
Swainson’s thrushes significantly changed the minimum frequency, bandwidth, and duration, of their songs upon exposure to the experimental traffic noise playback. The mean ± standard error for each individual bird is shown for (A) minimum frequency, (B) maximum frequency (C) bandwidth, (D) peak frequency, (E) duration, and (F) song rate (songs/minute) of Swainson’s thrushes’ full songs during the three playback phases (pre, noise, and post). Each color represents an individual Swainson’s thrush with lines connecting their response in each playback phase. Some individuals only sang in one or two of the phases. There are no connecting lines for the individuals who vocalized in pre- and post-playback but not during.

We found that the lower frequency introductory notes were also heavily impacted by noise playback (Table 2). Swainson’s thrushes significantly increased the minimum (noise, p<0.001; post, p<0.001; Figure 7A), maximum (noise, p<0.001; post, p = 0.004; Figure 7B), and peak frequency (noise, p<0.001; post, p = 0.014; Figure 7D) of their intro notes during noise playback and afterwards. The increase of these vocalization characteristics was relatively stronger during noise playback compared to after playback. Additionally, this increase was similar for the minimum and maximum frequencies as they went up by 390.45 ± 28.82 Hz and 422.31 ± 40.33 Hz, respectively, during noise playback and 111.88 ± 26.68 Hz and 107.11 ± 37.22 Hz once the noise stopped playing. The intro notes’ bandwidth (noise, p = 0.359; post, p = 0.789; Figure 7C) showed no significant changes during or after noise playback. Duration showed a non-significant trend as well; however, during noise playback there was a visible and near-significant increase in the length of the intro note (noise, p = 0.051; post, p = 0.944; Figure 7E). The traffic noise playback explained a meaningful proportion of the variation in the minimum, maximum, and peak frequency (marginal R^2^ = 0.325, 0.244, 0.250, respectively) of intro notes while only moderately or poorly explaining that of bandwidth and duration (R^2^ = 0.004, 0.016, respectively). While there was some variation in introductory syllables across different individuals, most birds responded to noise playback in a similar fashion (Figure 8).

**Figure 7.**
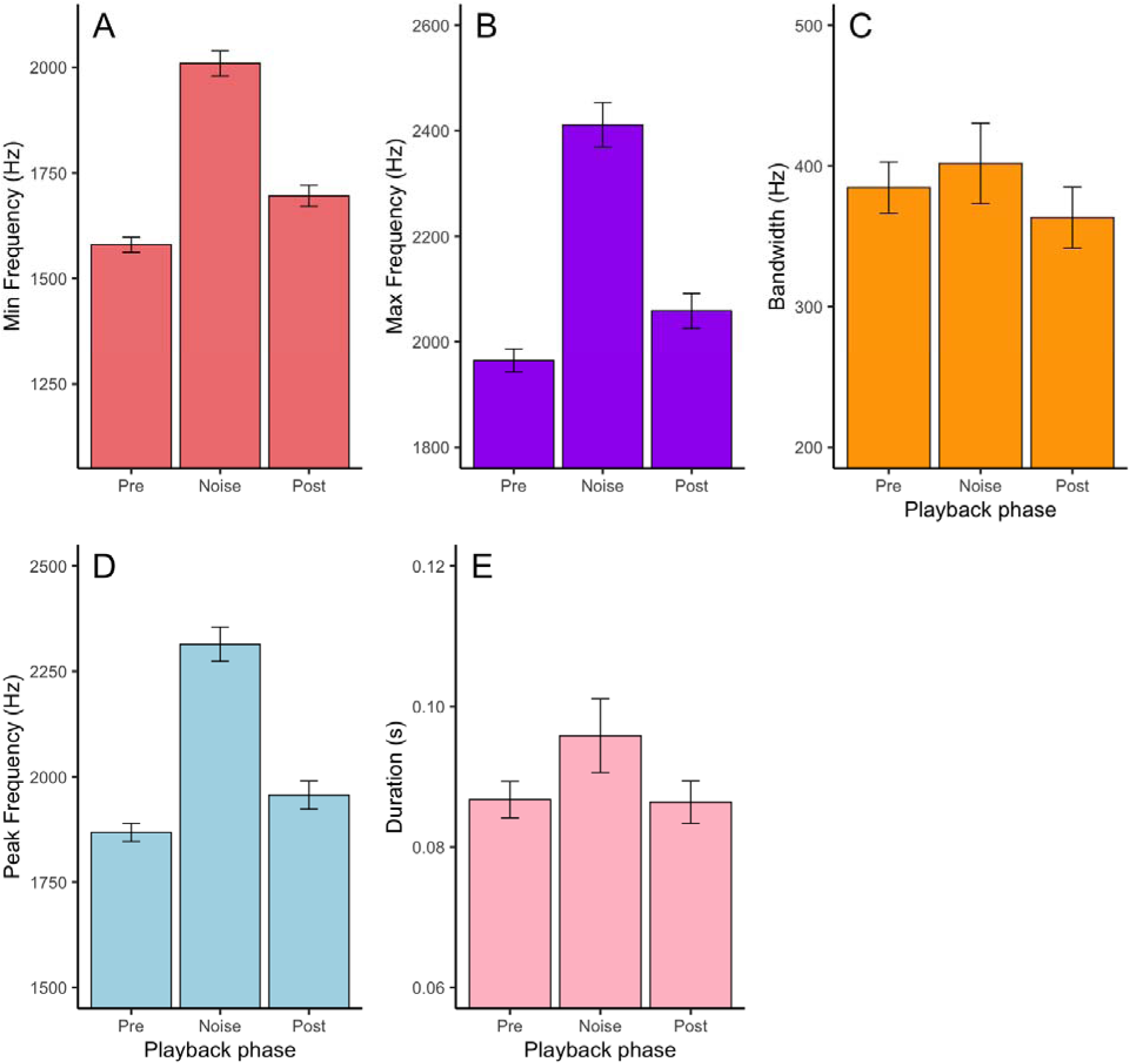
Swainson’s thrushes rapidly changed the minimum, maximum, and peak frequency of the intro notes of their songs upon exposure to the experimental traffic noise playback. Mean ± standard error are shown for the (A) minimum frequency, (B) maximum frequency (C) bandwidth, (D) peak frequency, and (E) duration of Swainson’s thrushes’ intro notes during the three playback phases (pre, noise, and post).

**Figure 8.**
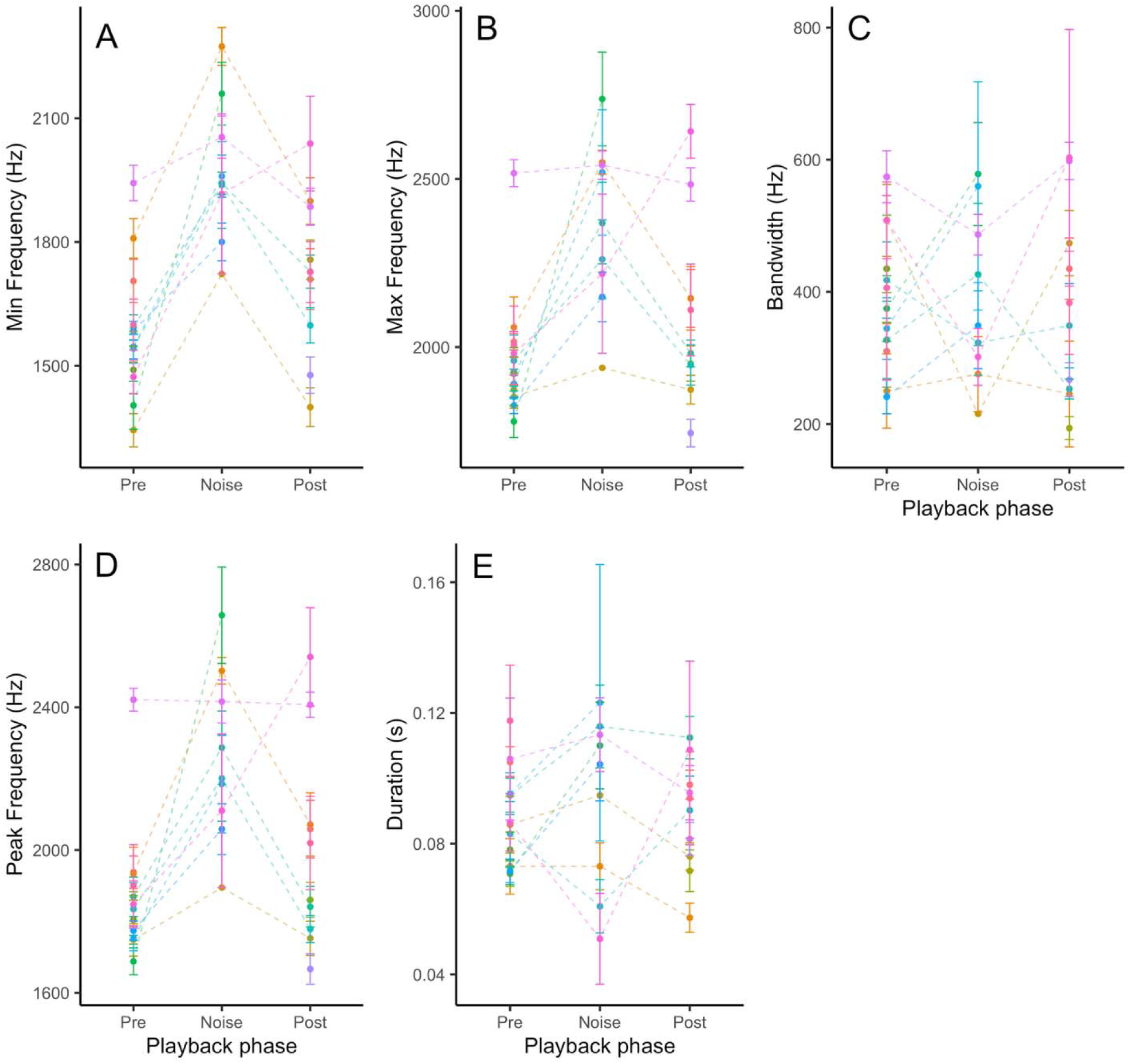
Swainson’s thrushes significantly changed the minimum, maximum, and peak frequency of the intro notes of their songs upon exposure to the experimental traffic noise playback. The mean ± standard error for each individual bird is shown for the (A) minimum frequency, (B) maximum frequency (C) bandwidth, (D) peak frequency, and (E) duration of Swainson’s thrushes’ intro notes during the three playback phases (pre, noise, and post). Each unique color represents an individual Swainson’s thrush with lines connecting their vocal response in each playback phase. Some individuals only vocalized in one or two of the phases. There are no connecting lines for the individuals who only vocalized during pre- and post-playback to avoid misinterpretation.

## DISCUSSION

We investigated whether Swainson’s thrushes vary their song structure and behavior in response to anthropogenic noise by analyzing frequency, duration, bandwidth, and song rate with respect to both population level variation across an ambient noise gradient and through short-term experimental exposure of individuals to traffic noise playback. Our results showed significant relationships between Swainson’s thrushes’ song structure and anthropogenic noise, both through the population level survey and individual level experimental data, indicating that they respond to short-term and long-term anthropogenic noise exposure by varying their singing behavior and song selection. The most consistent response to noise pollution was an increase in the minimum frequency of their songs. Birds also modified the maximum frequency, bandwidth, and duration of songs, but results vary between the individual and population levels. When examining the typically lower frequency introductory note of songs, we found changes in the maximum, minimum, and peak frequency at both the individual and population levels. Together, these results suggest that Swainson’s thrushes employ a variety of mechanisms to avoid vocal masking from low frequency traffic noise.

Our population level results show that Swainson’s thrushes residing in areas with higher levels of anthropogenic noise sang songs with a narrower bandwidth, characterized by increased minimum frequency and decreased maximum frequency. The increased minimum frequency was expected, as similar studies have shown that this is a common response used by a variety of different species to avoid masking by low frequency anthropogenic noise (e.g., Derryberry et al., 2016; Slabbekoorn et al., 2007). A finding unique to our study was the population-level decrease in maximum frequency. Swainson’s thrushes in higher noise areas may have been selectively singing songs within their repertoire that have smaller bandwidth (Halfwerk & Slabbekoorn, 2009), though it is not clear exactly how this change would be adaptive. At the individual level, Swainson’s thrushes faced with experimental traffic noise also increased their minimum song frequencies, but unlike at the population level, maximum frequencies showed no significant changes and in turn resulted in a less exaggerated reduction in bandwidth. Reduced song bandwidth may make vocalizations more audible over traffic noise, but it can also be detrimental to communication and mating by reducing song complexity and thus, potentially, attractiveness to mates (Winandy et al, 2021).

At the population level, Swainson’s thrushes showed no significant change in song duration. However, on the individual level, they increased song duration in response to noise playback. Previous studies have found conflicting results regarding changes to song duration, likely meaning that duration changes are species dependent and possibly influenced by other factors such as frequency of vocalization, habitat, and type of noise pollution (Francis et al., 2011; Bermudez-Cuamatzin et al., 2011). Similar studies that found increases in song duration propose that increasing song duration is an immediate strategy to compete with a novel noise (Brumm & Slater, 2006; LaZerte et al., 2017). Similarly, the immediate increase in Swainson’s thrush song duration shown by individuals confronted with experimental traffic noise could be an attempt to communicate over the noise, indicating short-term behavioral plasticity which would be unsustainable at longer periods of noise disturbance. Further work examining why Swainson’s thrushes did not demonstrate such plasticity on a population level may elucidate the long-term effects of anthropogenic noise on songbirds.

Results from both the population level and individual level of the study showed an increase in minimum, maximum, and peak frequencies in the introductory notes. These results were expected, as intro notes are the lowest frequency in the ascending progression of Swainson’s thrush songs (Dobson and Lemon, 1977). These low frequencies have a larger degree of spectral overlap with traffic noise, making them the most vulnerable to masking. This result is consistent with other studies which showed an increase in frequency of the introductory or lowest notes in low-frequency songs (Parris & Schneider, 2009).

Results for song rate were not significant in this study, though we did observe a general trend of decreased singing during experimental playback compared to the pre- and post-playback periods. It is possible that Swainson’s thrushes are assessing noise level variations and waiting for a quieter opportunity to sing. A study investigating the relationship between daytime noise levels and European robins’ (*Erithacus rubecula*) singing behavior found that robins sang nocturnally in areas with higher daytime noise levels, likely to take advantage of quieter conditions or avoid the competition from anthropogenic noise (Fuller, 2007). Further research could investigate if Swainson’s thrushes shifted from songs to calls during noise exposure, a pattern that has been observed in other species (LaZerte et al., 2017).

While Swainson’s thrushes are still a relatively common species in northern forests, their population has declined in many places, likely in part due to the loss of important habitat such as mature forests and riparian zones (Mack and Yong, 2000). One study found that out of 244 Swainson’s thrushes nesting in BC, Canada, 84% nested in undisturbed forest area (Campbell et al., 1997). We observed a similar trend during our data collection, finding more vocalizing males in relatively quieter areas compared to very loud roadside habitats (Pers. Obs.). Even with their apparent ability to immediately adjust song structure, Swainson’s thrushes may still seek out pristine habitats and avoid noisier locations, implying that noise impacts their assessment of habitat quality (Ware et al., 2015). The impacts of vocal masking have many detrimental effects on avian communication, and recent literature points to potential costs of changing song structure in response to noise including impaired territorial interactions between males, increased risk of predation or parasitism, and reduced clutch sizes in noisy nesting areas (Read et al., 2014; Derryberry et al., 2016; Francis et al., 2011). Investigating survival and reproduction rates of Swainson’s thrushes in environments that range in noise level would give insight into longer term impacts of anthropogenic noise.

Together our findings clearly show that Swainson’s thrushes’ song structure is influenced by background noise levels, with songs varying in minimum, maximum, and peak frequencies as well as duration in areas with higher noise levels. Additionally, they utilize a variety of mechanisms to change their song structure and singing behavior in real time in response to changes in anthropogenic noise levels, indicating the ability to adjust to short-term bursts of traffic noise. The latter shows that Swainson’s thrushes are among the species that have the vocal plasticity to rapidly assess variation in noise levels and adjust their vocalizations. Together, our results add to the growing body of research demonstrating that anthropogenic noise impacts avian communication and highlighting the diverse strategies that animals use to attempt to overcome these challenges for communication.

## ACKNOWLEDGEMENTS

This research was funded by the National Science Foundation (IOS-2207395) and Western Washington University. All work with animals was approved by the Western Washington University IACUC (#23-711).

## REFERENCES

Bates, D., Mächler, M., Bolker, B., & Walker, S. (2015). Fitting linear mixed-effects models using lme4. Journal of Statistical Software, 67, 1–48. 10.18637/jss.v067.i01

Bermúdez-Cuamatzin, E., Ríos-Chelén, A. A., Gil, D., & Garcia, C. M. (2011). Experimental evidence for real-time song frequency shift in response to urban noise in a passerine bird. Biology Letters, 7(1), 36–38. 10.1098/rsbl.2010.0437

Blackburn, G., Dutour, M., Ashton, B. J., Thornton, A., & Ridley, A. R. (2024). Anthropogenic noise affects vocalisation properties of the territorial song of western Australian magpies. BioRxiv. 10.1101/2024.02.15.578887

Brumm, H., & Zollinger, S. A. (2011). The evolution of the lombard effect: 100 years of psychoacoustic research. Behaviour, 148(11/13), 1173–1198.

Brumm, H., & Zollinger, S. A. (2013). Avian vocal production in noise. Animal Communication and Noise, 187-227. 10.1007/978-3-642-41494-7_7

Brumm, H., & Slater, P.J.B. (2006). Ambient noise, motor fatigue, and serial redundancy in chaffinch song. Behavioral Ecology and Sociobiology, 60(4), 475–81. 10.1007/s00265-006-0188-y.

Bruintjes, R., & Radford, A. N. (2013). Context-dependent impacts of anthropogenic noise on individual and social behaviour in a cooperatively breeding fish. Animal Behaviour, 85(6), 1343–1349. 10.1016/j.anbehav.2013.03.025

Campbell, R.W., Dawe, N. K., McTaggart-Cowan, I., Cooper, J. M., Kaiser, G. W., McNall, M. C. E., Smith, G. E. J. (1997). The birds of British Columbia, vol. 3: flycatchers through vireos. University of Chicago Press.

Charif, R. A., Waack, A. M., & Strickman, L. M. (2010). Raven Pro 1.4 user’s manual. Cornell Lab of Ornithology.

Collins, S. (2004). Vocal fighting and flirting: The functions of birdsong. In Nature’s Music (pp. 39–79). Elsevier.

C.W. Dobson, & Lemon, R.E. (1977). Markovian versus rhomboidal patterning in the song of Swainson’s thrush. Behaviour, 62(3/4), 277–297. http://www.jstor.org/stable/4533841

Derryberry, E. P., Phillips, J. N., Derryberry, G. E., Blum, M. J., & Luther, D. (2020). Singing in a silent spring: Birds respond to a half-century soundscape reversion during the COVID-19 shutdown. Science, 370(6516), 575–579. 10.1126/science.abd5777

Dowling, J. L., Luther, D. A., & Marra, P. P. (2012). Comparative effects of urban development and anthropogenic noise on bird songs. Behavioral Ecology, 23(1), 201–209. 10.1093/beheco/arr176

Esri. (2025) *Light Gray Reference* [Basemap layer]. https://basemaps.arcgis.com/arcgis/rest/services/World_Basemap_v2/VectorTileServer

Esri. (2022). *ArcGIS Pro (Version 3.0.0)* [Computer software]. Environmental Systems Research Institute. https://www.esri.com/en-us/arcgis/products/arcgis-pro/

Foote, A.D., Osborne, R.W. & Hoelzel, A.R., (2004). Whale-call response to masking boat noise. Nature, 428(6986), 910–910. 10.1038/428910a

Francis, C. D., Ortega, C. P., & Cruz, A. (2011). Different behavioural responses to anthropogenic noise by two closely related passerine birds. Biology Letters, 7(6), 850– 852. 10.1098/rsbl.2011.0359

Fuller, R. A., Warren, P. H., & Gaston, K. J. (2007). Daytime noise predicts nocturnal singing in urban robins. Biology Letters, 3(4), 368–370. 10.1098/rsbl.2007.0134

Gill, S. A., Job, J. R., Myers, K., Naghshineh, K., & Vonhof, M. J. (2015). Toward a broader characterization of anthropogenic noise and its effects on wildlife. Behavioral Ecology, 26(2), 328–333. 10.1093/beheco/aru219

Halfwerk, W., & Slabbekoorn, H. (2009). A behavioural mechanism explaining noise-dependent frequency use in urban birdsong. Animal Behaviour, 78(6), 1301–1307. 10.1016/j.anbehav.2009.09.015

Hohl, L. (2025). Galapagos yellow warblers differ in behavioural plasticity in response to traffic noise depending on proximity to road. Animal Behaviour, 222. 10.1016/j.anbehav.2025.123119

Hou, Z., Zhang, C., Li, L., Gao, B., Wu, R., Pei, N., & Liu, Y. Anthropogenic noise and habitat structure shaping dominant frequency of bird sounds along urban gradients. iScience, 27(2), 109056. 10.1016/j.isci.2024.109056.

Injaian, A. S., Taff, C. C., & Patricelli, G. L. (2018a). Experimental anthropogenic noise impacts avian parental behaviour, nestling growth and nestling oxidative stress. Animal Behaviour, 136, 31–39. 10.1016/j.anbehav.2017.12.003

Jacqmin-Gadda, H., Sibillot, S., Proust, C., Molina, J. M., & Thiébaut, R. (2007). Robustness of the linear mixed model to misspecified error distribution. Computational Statistics & Data Analysis, 51(10), 5142–5154. 10.1016/j.csda.2006.05.021

Job, J. R., Kohler, S. L., & Gill, S. A. (2016). Song adjustments by an open habitat bird to anthropogenic noise, urban structure, and vegetation. Behavioral Ecology, 27(6), 1734–1744. 10.1093/beheco/arw105

Kern, J.M. and Radford, A.N., (2016). Anthropogenic noise disrupts use of vocal information about predation risk. Environmental Pollution, 218, 988–995. 10.1016/j.envpol.2016.08.049

Kuitunen, M. T., Viljanen, J., Rossi, E., & Stenroos, A. (2003). Impact of busy roads on breeding success in pied flycatchers *Ficedula hypoleucaa*. Environmental Management, 31, 0079–0085. 10.1007/s00267-002-2694-7

LaZerte, S. E., Otter, K. A., & Slabbekoorn, H. (2017). Mountain chickadees adjust songs, calls and chorus composition with increasing ambient and experimental anthropogenic noise. Urban Ecosystems, 20, 989–1000. 10.1007/s11252-017-0652-7

Laiolo, P. (2010). The emerging significance of bioacoustics in animal species conservation. Biological Conservation, 143(7), 1635–1645. 10.1016/j.biocon.2010.03.025

Lüdecke, D., Ben-Shachar, M. S., Patil, I., Waggoner, P., & Makowski, D. (2021). Performance: an R package for assessment, comparison and testing of statistical models. Journal of Open Source Software, 6(60). 10.21105/joss.03139

Luke, S. G. (2017). Evaluating significance in linear mixed-effects models in R. Behavior Research Methods, 49, 1494–1502. 10.3758/s13428-016-0809-y

Luther, D. A., Phillips, J., & Derryberry, E. P. (2016). Not so sexy in the city: urban birds adjust songs to noise but compromise vocal performance. Behavioral Ecology, 27(1), 332–340. 10.1093/beheco/arv162

Montague, M. J., Danek-Gontard, M., & Kunc, H. P. (2013). Phenotypic plasticity affects the response of a sexually selected trait to anthropogenic noise. Behavioral Ecology, 24(2), 343–348. 10.1093/beheco/ars169

Mack, D.E., & Yong, W. (2000). Swainson’s thrush (*Catharus ustulatus)*. The Birds of North America, (540), 32.

Mulholland, T. I., Ferraro, D. M., Boland, K. C., Ivey, K. N., Le, M. L, LaRiccia, C. A., Vigianelli, J. M., Francis, C. D. (2018) Effects of experimental anthropogenic noise exposure on the reproductive success of secondary cavity nesting birds. Integrative and Comparative Biology, 58(5), 967–976. 10.1093/icb/icy079

Nakagawa, S., Johnson, P. C., & Schielzeth, H. (2017). The coefficient of determination R 2 and intra-class correlation coefficient from generalized linear mixed-effects models revisited and expanded. Journal of the Royal Society Interface, 14(134), 20170213. 10.1098/rsif.2017.0213

Nemeth, E., Pieretti, N., Zollinger, S. A., Geberzahn, N., Partecke, J., Miranda, A. C., & Brumm, H. (2013). Bird song and anthropogenic noise: Vocal constraints may explain why birds sing higher-frequency songs in cities. Proceedings of the Royal Society B: Biological Sciences, 280(1754), 20122798. 10.1098/rspb.2012.2798.

Parks, S. E., Johnson, M., Nowacek, D., & Tyack, P. L. (2011). Individual right whales call louder in increased environmental noise. Biology Letters, 7(1), 33–35. 10.1098/rsbl.2010.0451

Parris, K. M., & Schneider, A. (2009). Impacts of traffic noise and traffic volume on birds of roadside habitats. Ecology and Society, 14(1). https://www.jstore.org/stable/26268029

Read, J., Jones, G., Radford, A. N. (2013). Fitness costs as well as benefits are important when considering responses to anthropogenic noise. Behavioral Ecology, 25(1), 4–7. 10.1093/beheco/art102

Rhodes, M.L., Ryder, T.B., Evans, B.S., To, J.C., Neslund, E., Will, C., O’Brien, L.E. and Moseley, D.L. (2023). The effects of anthropogenic noise and urban habitats on song structure in a vocal mimic; the gray catbird (D*umetella carolinensiss*) sings higher frequencies in noisier habitats. Frontiers in Ecology and Evolution, 11, 1252632. 10.3389/fevo.2023.1252632

Schielzeth, H., Dingemanse, N. J., Nakagawa, S., Westneat, D. F., Allegue, H., Teplitsky, C., … & Araya Ajoy, Y. G. (2020). Robustness of linear mixed effects models to violations of distributional assumptions. Methods in ecology and evolution, 11(9), 1141–1152. 10.1111/2041-210X.13434

Slabbekoorn, H., & Ripmeester, A. A. P. (2008). Birdsong and anthropogenic noise: implications and applications for conservation. Molecular Ecology, 17(1), 72–83. 10.1111/j.1365-294X.2007.03487.x.

Slabbekoorn, H., & Den Boer-Visser, A. (2006). Cities change the songs of birds. Current Biology, 16(23), 2326–2331. 10.1016/j.cub.2006.10.008

Sordello, R., Ratel, O., Flamerie De Lachapelle, F., Leger, C., Dambry, A., & Vanpeene, S. (2020) Evidence of the impact of noise pollution on biodiversity: a systematic map. Environmental Evidence, 9, Article 20. 10.1186/s13750-020-00202-y

Templeton, C. N., Zollinger, S. A., & Brumm, H. (2016). Traffic noise drowns out great tit alarm calls. Current Biology, 26(22), R1173–R1174. 10.1016/j.cub.2016.09.058

Wang, W., Gao, H., Li, C., Deng, Y., Zhou, D., Li, Y., Zhou, W., Luo, B., Liang, H., Liu, W., Wu, P., Jing, W., & Feng, J. (2022). Airport noise disturbs foraging behavior of Japanese pipistrelle bats. Ecology and Evolution, 12(6), e8976. 10.1002/ece3.8976

Ware, H., McClure, C. J. W., Carlisle, J., & Barber, J. R. (2015). A phantom road experiment reveals traffic noise is an invisible source of habitat degradation. Proceedings of the National Academy of Sciences of the United States of America112(39)12105–12109. 10.1073/pnas.1504710112

Waser, P. M., & Waser, M. S. (1977). Experimental studies of primate vocalization: Specializations for long distance propagation. Zeitschrift für Tierpsychologie, 43(3), 239–263. 10.1111/j.1439-0310.1977.tb00073.x

Wiley, R. H., Richards, D. G. (1978). Physical constraints on acoustic communication in the atmosphere: Implications for the evolution of animal vocalizations. Bevavioural Ecology and Sociobiology. 3, 69–94. 10.1007/BF00300047

Winandy, G. S. M., Félix, R. P., Sacramento, R. A., Mascarenhas, R., Batalha-Filho, H., Japyassú, H. F., Izar, P., & Slabbekoorn, H. (2021). Urban noise restricts song frequency bandwidth and syllable diversity in bananaquits: Increasing audibility at the expense of signal quality. Frontiers in Ecology and Evolution, 9. 10.3389/fevo.2021.570420

Wood, W. E., & Yezerinac, S. M. (2006). Song sparrow (*Melospiza melodia*) song varies with urban noise. The Auk, 123(3), 650–659. 10.1093/auk/123.3.650

